# Understanding the high-order network plasticity mechanisms of ultrasound neuromodulation

**DOI:** 10.1101/2025.01.11.632528

**Authors:** Marilyn Gatica, Cyril Atkinson-Clement, Carlos Coronel-Oliveros, Mohammad Alkhawashki, Pedro A. M. Mediano, Enzo Tagliazucchi, Fernando E. Rosas, Marcus Kaiser, Giovanni Petri

## Abstract

Transcranial ultrasound stimulation (TUS) is an emerging non-invasive neuromodulation technique, offering a potential alternative to pharmacological treatments for psychiatric and neurological disorders. While functional analysis has been instrumental in characterizing TUS effects, understanding the underlying mechanisms remains a challenge. Here, we developed a whole-brain model to represent functional changes as measured by fMRI, enabling us to investigate how TUS-induced effects propagate throughout the brain with increasing stimulus intensity. We implemented two mechanisms: one based on anatomical distance and another on broadcasting dynamics, to explore plasticity-driven changes in specific brain regions. Finally, we highlighted the role of higher-order functional interactions in localizing spatial effects of off-line TUS at two target areas—the right thalamus and inferior frontal cortex—revealing distinct patterns of functional reorganization. This work lays the foundation for mechanistic insights and predictive models of TUS, advancing its potential clinical applications.

**Significance Statement:** Transcranial ultrasound stimulation (TUS) offers a non-invasive approach to modulating brain activity, holding promise for treating psychiatric and neurological disorders. Despite its potential, the mechanisms underlying its effects remain poorly understood. By integrating human fMRI data with whole-brain computational models, we identified how high-order functional interactions localize and propagate TUS-induced effects from local to global brain scales. This work introduces two mechanisms—distance-based propagation and diffusion-like broadcasting—that predict functional plasticity changes, providing a foundation for understanding and optimizing the biological and cognitive outcomes of TUS. Our findings offer critical insights into the dynamics of neuromodulation, bridging experimental results and clinical applications.

## I. INTRODUCTION

Non-invasive neuromodulation techniques have been gaining ground as an alternative to pharmacological interventions for the treatment of psychiatric and neurological conditions [1–3]. Existing tools can modulate neuronal firing rates via techniques including transcranial direct current stimulation, transcranial magnetic stimulation, and low-intensity transcranial ultrasound stimulation (TUS) — with the latter being capable of reaching deep brain areas with high spatial resolution [4–6]. Still, while TUS is a promising neuromodulation technique, many challenges remain in understanding its underlying mechanisms, for example how it translates into either stimulation or inhibition of neural activity [6–9]. Challenges in advancing the understanding of TUS effects include: (i) disentangling the spatially widespread changes generated by stimulus-induced plasticity, (ii) moving from population-level to individual-level descriptions, which are essential to designing personalized therapies, and (iii) predicting TUS effects via biologically realistic mechanisms. Here, we address these challenges by joining high-order interdependencies [10], communication models [11], and whole-brain modeling [12, 13]. Our aim is to identify robust informational markers to assess the alterations in brain function induced by stimulation, and to uncover their underlying biophysical mechanisms.

There are multiple ongoing research efforts trying to unravel the widespread changes generated by TUS. At the cellular level, the interaction of acoustic waves with the neuronal membrane in TUS has been investigated in terms of the activation of mechanosensitive ion channels or astrocytic TRPA1 [14–16], GABA inhibition [17] or cavitation [18, 19]. These mechanisms are related to synaptic plasticity processes, such as Long-Term Potentiation (LTP) and Long-Term Depression (LTD), by modulating neuronal excitability and neurotransmitter release [20]. At a global level, studies on functional connectivity have shown the impact of stimulus-induced plasticity at the population-level, revealing both increases and decreases in connectivity [21–24]. Additionally, high-order informational dependencies (HOI) have been used to characterize how TUS reorganizes the brain at the individual-level [25]. The core advancement of these methods with respect to traditional functional connectivity lies in their capacity to encode redundancy and synergy among signals [26–28]. For a simple example of the additional information encoded by these quantities, consider cooking. Individual ingredients might not provide a notable impact on texture, smell, and taste. However, when combined in a recipe, they create a different and –hopefully– memorable experience by working synergistically. Conversely, using multiple ingredients with similar texture or flavor would result redundant, because the same information is present among multiple ingredients.

After gaining insight into the functional changes induced by stimulation, we can use this knowledge to develop models that explain the plasticity-driven effects. This requires two elements: communication models, and wholebrain models. The former, communication models, are needed to describe how stimuli propagate across anatomically connected regions [11, 29–31]. In this context, navigation frameworks, such as shortest path length, have been widely applied to characterize neural communication [32–34]. However, diffusion models have recently demonstrated greater predictive power than efficiency-based approaches in explaining functional effects [11, 35], a notable example of this being recent results on the propagation of focal electrical stimulation in intracranial EEG recordings of drug-resistant epilepsy participants,[36]. Whole-brain models involved instead the integration of structural connectivity and neuronal dynamics [13, 37– 40] to enable the testing of mechanistic hypotheses, including biophysically inspired ones [12, 41].

Here, by combining these elements, we aim to elucidate: (i) to what extent high-order functional interactions to localize TUS-induced spatial effects; (ii) how widespread is the propagation of a stimulus across the brain when its magnitude increases; and (iii) which network communication model of the *TUS-induced plasticity* better explains mechanistically the *functional changes* induced by the stimulation. To address these questions, we analyze fMRI data of human subjects stimulated at two different targets: the right inferior frontal cortex and the right thalamus. We show that the TUS of each induces a specific signature of spatially widespread redundant and synergistic changes. Specifically, for the inferior frontal cortex stimulation (TUS-IFC), we observe effects in the frontal and basal ganglia areas, while for the thalamic stimluation (TUS-Thal), effects are prominent in the cingulate, temporal, and basal ganglia regions. Additionally, we find that communication models based on network communicability and distance are more reliable predictors of high-order functional changes than other communication models for both stimuli. Lastly, using the two most informative communication models for the plasticity, we develop a whole-brain model, reproducing the spreading of the stimulation throughout the brain as the stimulus intensity increases.

## II. RESULTS

We analyzed changes in redundancy and synergy in fMRI data following TUS. The participants underwent an initial fMRI session lasting approximately 14 minutes without stimulation (control, N = 22). On a separate day, they received 80 seconds of TUS, with participants receiving stimulation targeted at either the right inferior frontal cortex (TUS-IFC, N = 11) or the right thalamus (TUS-Thal, N = 11) (Fig. 1A), followed by an fMRI scan lasting around 42 minutes.

**FIG. 1.**
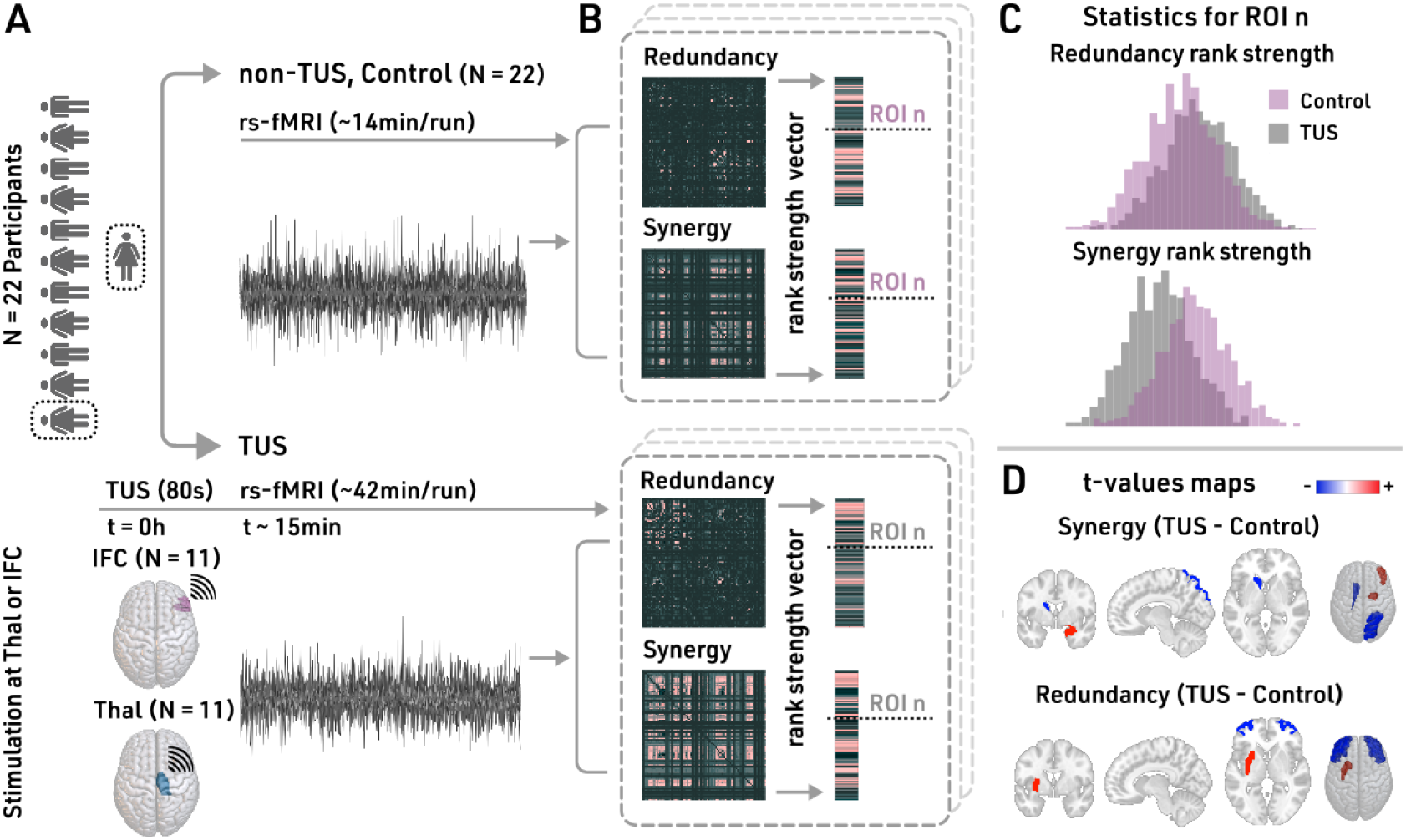
Methodology and absolute changes in redundancy and synergy after TUS. **A**. Twenty two subjects participated in the experiment, *N* = 22 controls (non-TUS), *N* = 11 IFC-TUS, and *N* = 11 Thal-TUS. **B**. We quantified the redundancy and synergy matrices, computing their median vector across rows and their ranked version, named medianredundancy-rank or median-synergy-rank. **C**. For each ROI, we compared each median-HOI-rank/absolute distribution (dotted line in **B**) between the control and TUS. **D**. We reported the t-values for the absolute changes, representing a shift to an increase (red) or decrease (blue) HOI interactions.

We calculated redundancy and synergy (Fig. 1B) for each pair of time series using the Integrated Information Decomposition method [28]. This approach decomposes the information of two variables measured at two consecutive time points into redundant, synergistic, and unique components that characterize specific dynamical patterns. Following previous work [42], our analysis focused only on the redundant and synergistic interactions, neglecting the unique information component (see *Methods* for further details). To assess the relative relevance of every brain region in synergistic or redundant interactions, we ranked regions by their redundancy-strength (median-redundancy-rank) and synergy-strength (median-synergy-rank).

### Transcranial ultrasound stimulation alters HOI, revealing spatial localization

To observe local functional changes due to stimulation, we compared the distributions of the redundancy (median-redundancy-rank) and synergy (median-synergy-rank) (Fig. 1B) between the control group (non-TUS) and each TUS condition for each brain area (Fig. 1C-D).

For TUS-IFC, we find alterations in the redundancy in frontal regions (parsorbitalis, right rostral middle frontal, right caudal middle frontal, and right paracentral), and basal ganglia areas (accumbens and caudate) (Fig. 2A, top row). Statistics (t-values and p-values) for these findings are reported in Supplementary Table 1. Results also show changes in synergy at frontal (rostral middle frontal), parietal (supramarginal), temporal (temporal pole, entorhinal), and basal ganglia regions (putamen) (Supplementary Table 1) (Fig. 2A, bottom row). In contrast, after TUS-Thal, the redundancy (Fig. 2B, top row) changed at frontal (parstriangularis, lateral orbitofrontal), temporal (superior temporal, middle temporal), basal ganglia (accumbens), cingulate (posterior cingulate), and occipital (lateral occipital). The synergy showed differences at the cingulate (rostral anterior cingulate), temporal (entorhinal), and basal ganglia areas (pallidum and thalamus). We report the statistics (*t*-values and *p*-values) in Supplementary Table 1.

**FIG. 2.**
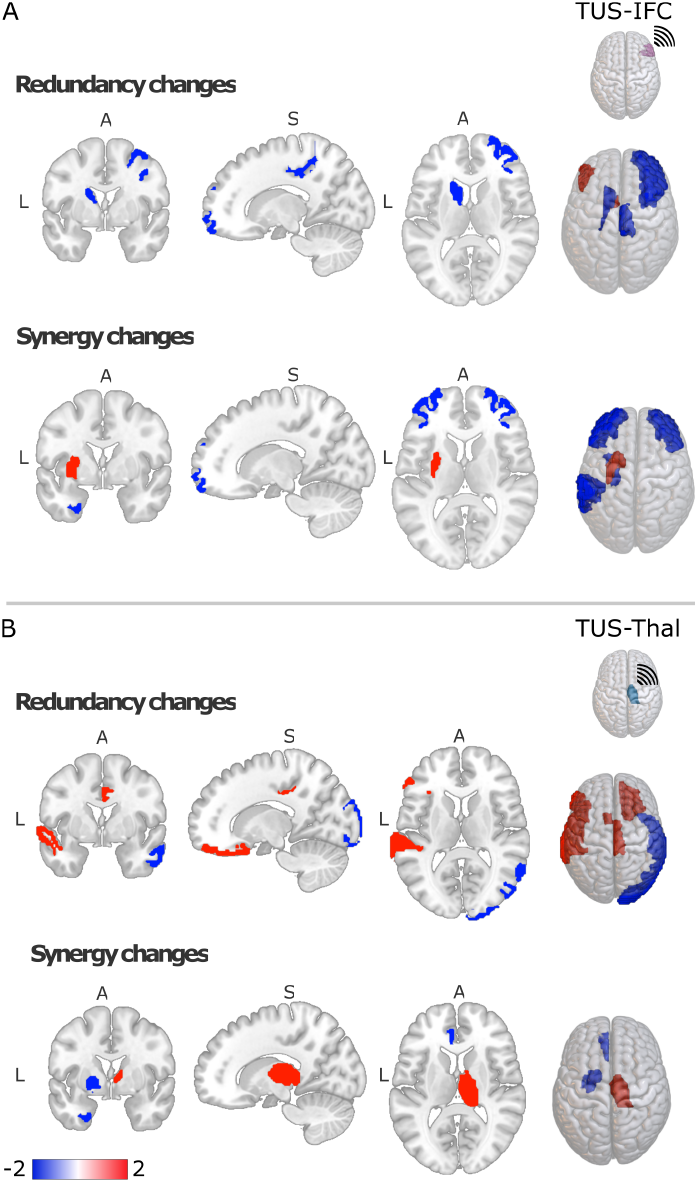
Relative changes in redundancy and synergy after TUS. **A**. Top row: median-redundancy-rank distribution changes after the TUS-IFC. Bottom row: Median-synergy-rank distribution changes after the TUS-IFC. The blue represents a region decreasing the HOI after TUS, wheres the red color describes the increase. **B**. Similar to A, when the target is the thalamus (TUS-Thal). We reported the t-values corrected by a *N* = 1000 permutation test in all the comparisons.

In conclusion, the functional effects induced by TUS vary significantly depending on the stimulation target, leading to distinct spatial patterns. Overall, after TUS-IFC redundancy and synergy show changes in the frontal and basal ganglia areas, with synergy additionally extending into the temporal and parietal lobes. When focusing on the frontal areas, the redundancy presented lateralized changes, mainly decreasing in the right hemisphere, particularly in the rostral and caudal middle frontal regions and the paracentral lobule. In contrast, after TUS-Thal, both quantities presented spatially widespread functional changes in the basal ganglia, temporal, and cingulate regions, extending redundancy into the frontal and occipital lobes. Interestingly, only the synergy showed an increase in the right thalamus, which was the targeted area for stimulation.

### Distance and communicability predict changes in HOI

After characterizing the local effects induced by TUS, we aim to determine which communication models best explain the global HOI effects produced by the stimulation. To approach this, we examine the associations of changes in redundancy and synergy with various models of stimulus propagation, based either on efficient navigation or on diffusion. In particular, following previous work (see [36] and *Methods* for further details), we adopt streamline length (distance) and shortest path efficiency (SPE) as proxies for efficiency, and search information (SI) and communicability (CMY) as a proxy for diffusion. To compute the associations between HOI changes and these models, we created eight “representative” vectors: one for each of the four connectivity models (distance, SPE, SI, CMY), and four representing redundancy and synergy changes for the two targets (TUS-IFC or TUS-Thal minus control), respectively. We describe these vectors as *representative*, because the models were computed using group-averaged properties (more specifically, average anatomical or functional matrices; see Methods for details), rather than individual measurements. Then, for each target, we correlated the vectors corresponding to the four models with those representing the changes in redundancy and synergy for that target, where by changes here we refer to the difference between the measurements after stimulation and those in the control condition (e.g. TUS-IFC minus control).

Surprisingly, we found two opposing patterns (Fig. 3). For TUS-IFC, redundancy alterations are negatively correlated with both network communicability (*r* = − 0.381, *p* < 0.001) and distance (*r* = − 0.569, *p <* 0.001), while synergy alterations are not significantly correlated with any of the two (Fig. 3A). For TUS-Thal, synergy alterations are positively correlated with both network communicability (*r* = 0.353, *p* = 0.001) and distance (*r* = 0.439, *p* < 0.001), while redundancy ones are not correlated with any of the two (Fig. 3B). All other communication models do not give any significant result, with exception of a negative correlation of synergy with shortest path efficiency for TUS-Thal (*r* = − 0.325, *p* = 0.003. Supplementary figure S1).

**FIG. 3.**
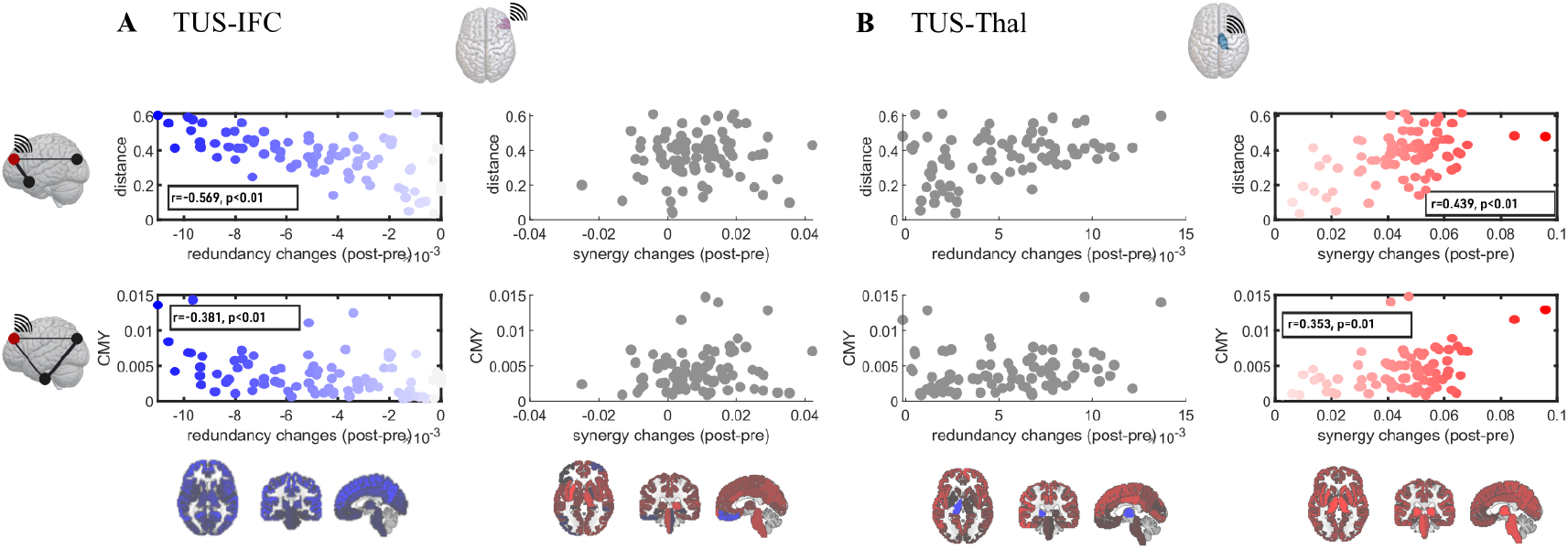
Whole-brain associations between structural models and observed changes in TUS. For TUS-IFC **A**. and TUS-Thal **B**., we computed the models and changes in HOI (after minus before) over a representative matrix (redundancy, synergy, distance, and the communicability, CMY) averaged across all participants. Within each subpanel, each row corresponds to a structural model (distance, top row; communicability model, CMY, bottom row), while each column corresponds to changes in informational quantities (redundancy, left column; synergy, left right column). The darker boxes represent the p-values lower than 0.05 after the Bonferroni correction, with the blue dots representing the redundant and red dots the synergistic changes. The grey colour dots represent the non-significant associations. For the other two models, see Supplementary figure S1.

In both cases, the largest changes in absolute value for both redundancy and synergy after stimulation were associated with longer distances and with regions (see Fig. 3A-B, top row). However, in this case too, the effects are opposing: for TUS-IFC we observe an overall decrease of redundancy with distance from the stimulation (Fig. 3A, top row); for TUS-Thal instead we find an overall increase in synergy with distance from the stimulation. Together these findings suggest the presence of a strong network effect in the TUS-induced plasticity, possibly mediated by the multiplicity of propagation paths between regions [43].

### Whole-brain model informed by distance or communicability heterogeneity explains changes in HOI

To test the hypothesis of a mechanistic link between the observed global effects of TUS on HOI and communication models encoding different notions of connectivity, we propose a whole-brain model which explicitly includes communicability and distance as mechanisms affecting redundancy and synergy. Specifically, we used a Hopf model of neural oscillators, in which the local dynamics of each node was simulated using the Stuart-Landau oscillator. For positive bifurcation parameter (*a* > 0), the model enters a limit cycle, and the system exhibits sustained oscillations. For negative bifurcation parameter (*a* < 0), the model has a stable fixed point, and thus the system will be dominated by noise. Finally, near the bifurcation point (*a* = 0), noise-driven and sustained oscillations coexist in time (Fig. 4A).

**FIG. 4.**
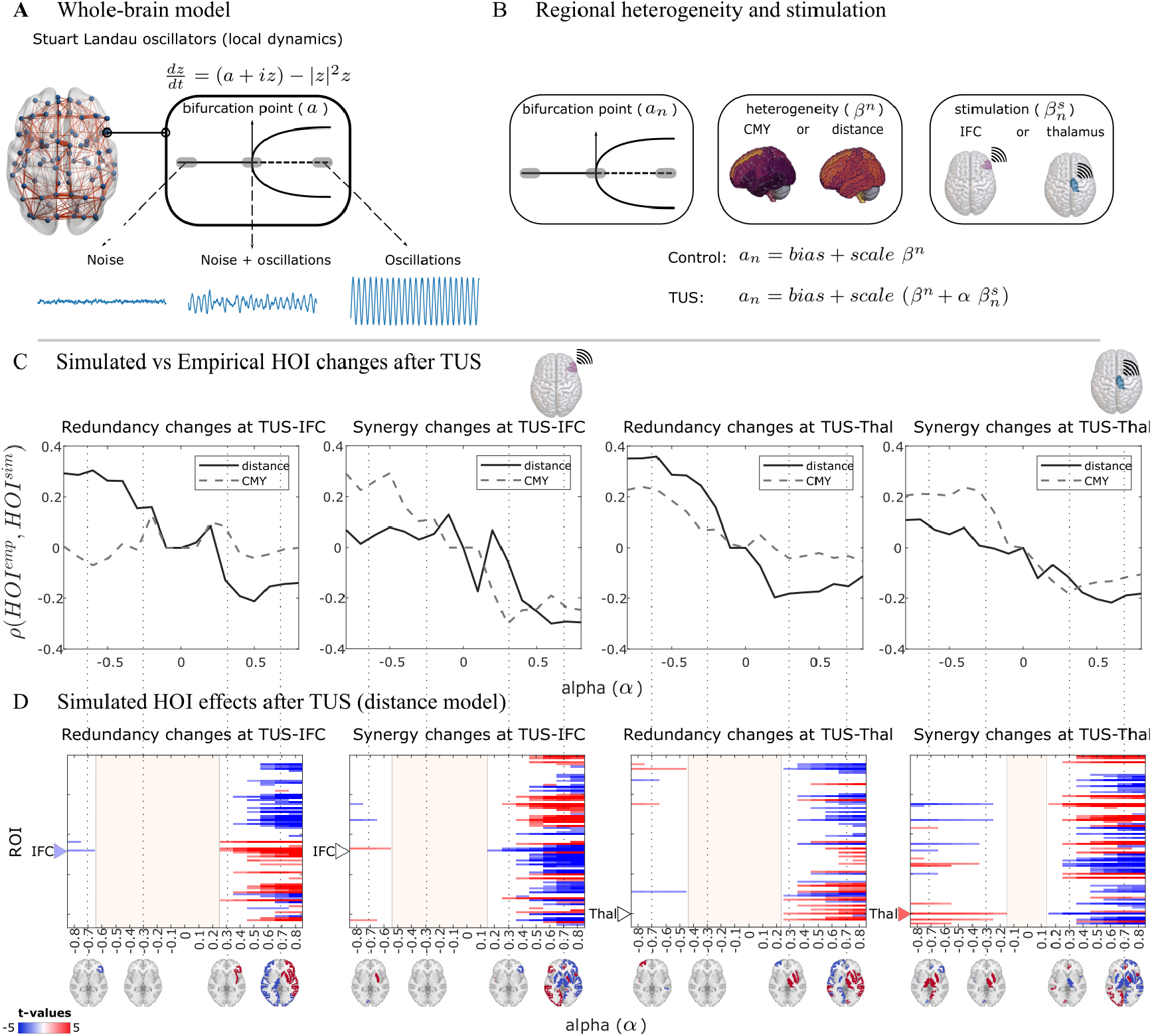
Whole-brain modeling predicts the propagation of TUS-induced plasticity from local to global scales. **A**. The local dynamics of each node were simulated using the Stuart-Landau oscillator, which, depending on the bifurcation parameter (a), can exhibit sustained oscillators (*a >* 0), noise (*a <* 0) or coexistence of noise-driven and sustained oscillations (*a* = 0). **B**. We inform heterogeneous models with communication models based on communicability (CMY) or distance (denoted as *β*^*n*^), and a stimulation modulated by the parameter *α* (denoted by 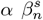) for each target. **C**. Redundancy and synergy fitting between the empirical and simulated data: The x-axis corresponds to different simulated intensities in the model (alpha), while the y-axis to the Spearman correlation between the empirical and simulated statistical differences (all the t-values in TUS minus control). Each column corresponds to the changes in redundancy and synergy for the two targets. Results for the model based on distance are shown with a solid line, those for the one based on the communicability model with a dashed line. **D**. Corrected *t*-values in the simulated data (TUS minus control) for the distance-based model (for the communicability model, see Fig S4). The columns are consistent with panel **C**. The brain plots illustrate the HOI changes, displaying significant t-values corrected using a permutation test with N = 1000 iterations. Colors indicate negative/positive changes with respect to no stimulation. Coloured triangles represent the stimulated target with significant t-values at negative alpha.

To include different notions of TUS propagation, we informed the model using communicability or distance (denoted as *β*^*n*^), and a stimulation strength (denoted by 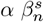) to quantify the changes in the TUS (Fig. 4B). Our approach involved a two-step fitting process as outlined in previous literature [44]. First, we performed a homogeneous fitting to determine the optimal global coupling (*G*). Then, we conducted a heterogeneous fitting to obtain the “bias” and “scale” parameters (Fig. 4B) to fit the control condition (see Supplementary Figure S2). In both cases, the goal was to achieve model simulations that reproduce as closely as possible the functional connectivity and mean brain synchrony (computed as the average Kuramoto order parameter, KOP) measured in data. Finally, after fitting the control condition, we simulated the effects of stimulation over a range of perturbation intensities (described by the parameter *α*).

Using the calibrated models, we examined the similarity between the empirical and simulated HOI effects of TUS, revealing three key findings. First, negative *α* values lead to better performance regardless of the target (Fig. 4C, *α* in the x-axis). Specifically, when *α* is negative, it results in larger positive correlations between the simulated and observed HOI values, whereas positive *α* values produced smaller, anti-correlated values. This suggests that the stimulation is likely disrupting the excitatory/inhibitory balance by increasing inhibition (noise), rather than by enhancing excitation (synchronization). Moreover, when *α* increased, the time series became progressively more synchronized, with the thalamus stimulation leading to faster system synchronization (see Supplementary Figure S3). Second, for both targets and negative *α*, redundancy changes were best reproduced by the distance-based model, while synergy changes were more accurately captured by the communicability model. Third, moving from strongly negative to strongly positive *α*, we consistently observed a local-to-global transition characterized by three main regimes: (i) only localized effects on a handful of regions for *α* ≪0, (ii) an intermediate regime in which no region shows significant alterations in their synergy/redundancy behaviours for *α* ∼ 0 (Fig. 4D, dashed red area), finally, (iii) global, delocalized effects across the whole brain for both redundancy and synergy for *α* ⪢ 0. Notably, in the simulated TUS-IFC condition, in the localized regime (*α* ≪ 0), we recovered the redundancy alterations in the right IFC (Fig. 4D, first panel, blue triangle) but not synergistic ones (Fig. 4D, second panel, white triangle). Conversely, for the TUS-Thal condition, only the synergistic alterations in the right thalamus were reproduced (Fig. 4D, fourth panel, red triangle).

## III. DISCUSSION

Our results revealed distinct patterns of functional reorganization following TUS, depending on the target. We obtained these findings combining three innovations: (i) the application of HOI to localize spatial effects induced by TUS in humans, extending previous studies in macaque data [21, 22, 25], (ii) the use of communication models as mechanisms to predict the functional plasticity-driven impact of TUS, and (iii) the development of a model to explain the mechanism of propagation of the effects of TUS when the stimulation intensity increases. We found that, for TUS-IFC, HOIs exhibited changes in frontal and basal ganglia areas, with the redundancy decreasing in the right frontal hemisphere, whereas, for TUS-Thal, changes were localized in the cingulate, temporal, and basal ganglia areas, with the synergy increasing in the stimulated right thalamus. Although the TUS protocol is designed to modulate neuronal activity by either increasing or decreasing it, studies have observed both increased and decreased functional connectivity [21–24], as well as higher-order interactions [25] in macaques following TUS. This variability may arise from a combination of LTP or LTD-like plasticity effects, resulting in heterogeneous outcomes [21, 22, 45]. We reported two possible mechanisms for plasticity induced by the stimulation to predict the functional changes after TUS. The global functional changes produced by TUS were associated with distance and communicability regardless of the targeted area. In turn, models based on distance and communicability outperformed models based on shortest path efficiency and search information. Altogether, our findings align with results from drug-resistant epilepsy participants [36], in reported communicability and search information —both diffusion processes— were found to be the best predictors for the propagation of focal electrical stimulation. Our results also align with previous research predicting functional connectivity patterns based on network communication, in which diffusion models were shown to outperform models based on shortest path length [35]. We believe that these findings provide valuable insights for modeling the effects of various types of stimulation and suggest potential avenues for further research and clinical applications [11].

We developed a whole-brain model explaining how the effects of TUS spread spatially throughout the brain as the stimulation intensity increases. In particular, depending on the intensity, the effects transitioned from a localized to a global regime, opening new paths for the exploration and prediction of changes in brain function. We also found that larger stimulation intensities led to quicker synchronization in the thalamus compared to the IFC. This can be explained by the thalamus’s more central role in system integration [46–49], which would increase its capacity to promote more synchronized interactions. Moreover, we implemented two possible mechanisms of TUS propagation, one based on distance and one broadcasting process that allowed us to test the plasticity-driven changes in some brain areas for different intensities. Similar to adjusting the bifurcation parameter in the Hopf model, biophysically-inspired models have shown a transition from noisy oscillations to sustained oscillations when the excitatory/inhibitory (E/I) balance is disrupted through increased inhibition or excitation, respectively [50–53]. In our findings, both models indicated that the stimulus is more likely to induce noise in the system rather than pure synchrony. This suggests that the stimulus tends to disrupt the E/I balance by enhancing inhibition rather than excitation. We stress that –although the mechanisms of TUS or the plasticityinduced chances are still a matter of debate [14–19]– which mechanism leads to either inhibitory or excitatory outcomes is even less understood [8, 17, 19, 54]. The theta-burst TUS protocol used in this study has been associated with an increase and decrease in neural activity [55], and our model could help clarify whether ultrasonic neuromodulation produces excitatory or inhibitory effects. The whole-brain models have been used on perturbations and psychiatric or neurological conditions, enabling to test mechanisms that can be used for predicting the outcomes of real experimental settings [37–40, 56]. Higher-order interactions have gained prominence in clinical applications for characterizing and predicting healthy aging [39, 57, 58], early development [59], neurological conditions [60, 61], and their associations with cognition [42] and consciousness [62]. Recently, HOI has been applied to transcranial ultrasound stimulation (TUS) in macaques, revealing different topological reorganizations depending on the stimulation target [25]. Here, we extended this understanding to healthy humans, demonstrating that spatial differences in response to TUS also rely on the stimulation target.

Before concluding, it is important to acknowledge the limitations of this study. First, our experiment included 22 participants. Future studies should build on this by incorporating a larger sample size to enhance the robustness of the results. Second, we proposed a mechanism for modifying regional excitability. While the effects of perturbing only the stimulated target have been observed, this presents an exciting avenue for future research, such as further exploring the impact of stimulating both the target and adjacent areas. Finally, while we used the minimum mutual information (MMI) redundancy function, as supported by previous literature demonstrating its clinical relevance in cognition [42], similar outcomes might be attainable with other redundancy functions [63]. Exploring alternative definitions [64, 65] could offer valuable opportunities for further investigation.

## IV. MATERIALS AND METHODS

### Partial Information Decomposition (PID)

Consider three random variables: two source variables *X*^*i*^ and *X*^*j*^, and a target variable *Y*. The Partial Information Decomposition (PID) [66] decomposes the total information provided by *X*^*i*^ and *X*^*j*^ about *Y*, given by Shannon’s mutual information *I*(*X*\*; *Y*), as follows:

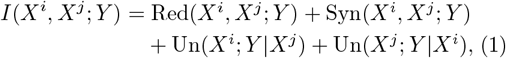

where Red(*X*^*i*^, *X*^*j*^; *Y*) represents the information provided by *X*^*i*^ and *X*^*j*^ about *Y* (redundancy), Syn(*X*^*i*^, *X*^*j*^; *Y*) denotes the information provided jointly by *X*^*i*^ and *X*^*j*^ about *Y* (synergy), Un(*X*^*i*^; *Y* |*X*^*j*^) is the unique information provided by *X*^*i*^ about *Y*, and Un(*X*^*j*^; *Y* |*X*^*i*^) is the information that is provided only by *X*^*j*^ about *Y*. The four terms of this decomposition are naturally structured into a lattice with nodes 𝒜= {{12}, {1}, {2}, {1} {2}}, corresponding to the synergistic, unique in source one, unique in source two, and redundant information, respectively. To compute these terms, we followed the minimum mutual information (MMI) PID decomposition for Gaussian systems [67], where the redundancy is computed as the minimum information between each source and the target, and the synergy refers to the additional information provided by the weaker source when the stronger source is known.

#### Integrated information decomposition (ΦID)

Consider the stochastic process of two random variables 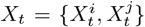 and denote the two variables in a current state *t*, by 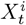 and 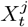, and the same two variables in a past state *t*− *τ*, by 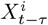, and, 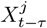. The integrated information decomposition (Φ*ID*) is the forward and backward decomposition of 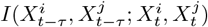, called the time delay mutual information, in redundant, synergistic and unique information [28]. The ΦID can be represented by the forward and backward interactions of the product 𝒜 × 𝒜, resulting in 16 distinct atoms: synergy to synergy, redundancy to redundancy, unique in source one to unique in source two (and backward), and redundancy to synergy, among others. Following previous work [42], our analyses focus on two specific atoms quantifying the temporal persistence of redundancy and synergy: persistent redundancy (redundancy that remains redundancy) and persistent synergy (synergy that remains synergy).

Note that while some approaches to assessing higher-order interactions involve three or more time series, we employed the “higher-order” concept here as we analyzed four random variables 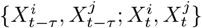, where (*X*^*i*^, *X*^*j*^) represent the two variables at the current state *t*, along with 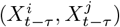 being their states at *t* − *τ*.

#### Synergy and redundancy rank framework

The ΦID was computed for all combinations of pairwise BOLD time series 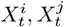, where *i* and *j* represent two different brain regions, with (*i, j*) ∈ 1, …, 84. This resulted in two symmetrical matrices capturing redundancy and synergy. We then calculated the median of each matrix to obtain two strength vectors (each 1×84) representing redundancy and synergy. Additionally, we derived rank strength vectors for both redundancy and synergy (each 1×84), ranking each region by its strength. Finally, we compared the absolute or rank strength values between the TUS (IFC-TUS or Thal-TUS) and the Control vector.

### Distance and communication models

*D* ∈ ℝ^84×84^ denotes the distance with *D*_*ij*_ being the average streamline length between two regions *i* and *j*, computed per subject. As a representative value of *distance*, we used an average across subjects normalized by the maximum 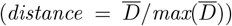. Similarly, *M* ∈ R^84*×*84^ denotes the structural connectivity, where *M*_*ij*_ is defined as the average number of streamlines connecting two brain regions *i* and *j*. We computed an average across subjects as the representative structural connectivity, denoted by 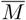, likewise normalized with real values between zero and one. Then, we computed the matrix of lengths 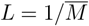, with *L*_*ij*_ the cost of communication between the regions *i* and *j*.

The *Shortest Path efficiency* is denoted by *SPE*, where (*SPE*_*ij*_) is computed as the inverse of the shortest path or geodesic connecting two nodes *i* and *j*. Given the sequence of regions *{i, u*, …, *v, j}*, such that 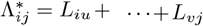 is the minimum transmission cost between *i* and *j*, then 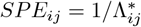 [68].

The *Search Information* is denoted by *SI* and quantifies the amount of information to bias a random walk into the shortest path *{i, u*, …, *v, j}*. The transition probability of traveling from *i* to *j* is computed as 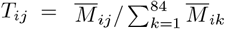. Therefore, the probability of a random walk to travel from *i* to *j* via the shortest path is Π_*ij*_ = *T*_*iu*_ *×* · · · *× T*_*vj*_. Finally, the search information (*SI*_*ij*_) is computed as *SI*_*ij*_ = − log_2_(Π_*ij*_) [35].

The *Communicability*, denoted by *CMY*, is a broadcasting process quantifying the redundant walks connecting two regions while penalizing the longer paths. Therefore, 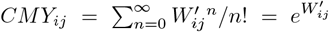, where 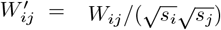, and 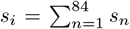 is the strength of region *i* [69, 70].

### Whole-brain model

We simulated the brain activity using a supercritical Hopf bifurcation model (Stuart-Landau oscillators) [37, 71]. The following ordinary differential equations [3] define the dynamic for each node *i*:

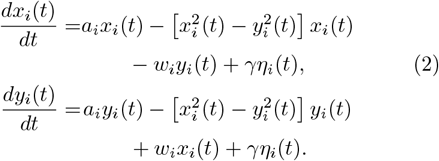

Where *y*(*t*) corresponds to the imaginary component, and the real component of the time series, *x*(*t*), simulated the BOLD-like signals. We set the oscillation frequency *f*_*i*_ = 0.05 Hz (*w*_*i*_ = 0.05 ×2 ×*π*) for overall nodes. When the bifurcation parameter *a* is positive (*a >* 0), the system exhibits a limit cycle, leading to sustained oscillations. If (*a <* 0), the system has a stable fixed point and is dominated by noise. Near the bifurcation point (*a* = 0), both noise-driven and sustained oscillations coexist over time. We used 84 nodes, parcellated using the Desikan-Killiany atlas (See Supplementary Tables S2 and S3 for details), including subcortical areas and structural connectivity matrices *M* ∈ R^84*×*84^ (19 matrices, as described in Supplementary information, three images were excluded due to excessive motion). The brain areas are coupled with the structural connectivity *M, G* = 0.16 represents the global coupling (a fitted parameter. Supplementary figure S2), and *η*_*i*_(*t*), with *γ* = 0.02 the standard deviation, the external Gaussian noise [3].

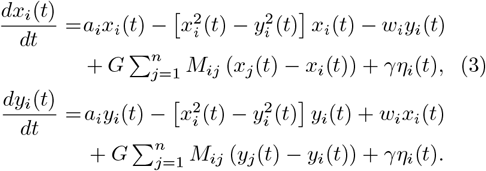

We proposed modifying the bifurcation parameter as a proxy of modulate neuronal activity. Biophysical-inspired models have revealed a switch from noisy oscillations to sustained oscillations (limit cycles) when disrupting the excitation/inhibition balance through excitation [50–53], analogous to increasing the bifurcation parameter in the Hopf model. Following previous literature, we incorporated heterogeneity in our model [44].

#### Control and TUS model

We simulated the control condition using the Hofp model with *a*_*i*_ = *bias* + *scale β*^*i*^, where *β* is the heterogeneous vector of distances or communicability. The parameters *scale* and *bias* were fitted as (*bias, scale*) = (−0.17, 0.24) in the distance model and (*bias, scale*) = (− 0.08, 0.21) in communicability (see Supplementary figure S2). Additionally, for TUS, 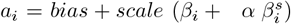, where 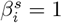 if *s* is the stimulated target and zero in other case.

We ran 1100s simulations with an integration step of 0.1s in the Euler-Maruyama integration scheme for each subject (19 structural connectivity matrices). The simulated time series were band-pass filtered between 0.001 and 0.01 Hz and removed the first and last 100s, resulting in 15-minute simulations.

### Statistical analysis

This study compared the redundancy and synergy of each control (non-TUS) versus each TUS experiment (IFC-TUS or Thal-TUS) using a t-stat (TUS minus control). We performed a 1.000 permutation two-samples t-test analysis per region in the empirical data and the simulations to find the statistically significant differences. Bonferroni corrected the correlations between redundancy and synergy changes with the communication models.

## Code availability

The data analysis was conducted using MATLAB version 2022b. The MATLAB code for quantifying synergy and redundancy from integrated information decomposition of time series, utilizing the Gaussian MMI solver, is available at https://doi.org/10.1038/s41593-022-01070-0 [42]. We computed the communication models for weighted networks using the Brain Connectivity Toolbox [72]. We used the Python code to simulate the Hopf model, freely available at: https://github.com/carlosmig/StarCraft-2-Modeling.git[71].

## ACKNOWLEDGMENTS

M.K. and C.A., were supported by the Engineering and Physical Sciences Research Council (EP/W004488/1, EP/X01925X/1 and EP/W035057/1). M.K. was also supported by the Guangci Professorship Program of Rui Jin Hospital (Shanghai Jiao Tong University).

