## Supplementary Material for "Understanding the high-order network plasticity mechanisms of ultrasound neuromodulation"

#### Participants

Twenty-two healthy, right-handed volunteers participated in this study. None had a history of neurological or psychiatric disorders (except for cases of depression considered remitted for at least one year) and were not taking any medications. The additional exclusion criteria included close relatives with a history of seizures, a predisposition for syncope, excessive hair that could interfere with transducer coupling, current or planned pregnancy, implanted metallic devices, skin diseases, claustrophobia, or anxiety related to MRI, and tattoos near the head. Participants were instructed to avoid recreational drugs for 48 hours before their visits and to limit alcohol consumption to no more than four units within the preceding 24 hours. The study adhered to the ethical standards of the Helsinki Declaration of 1978, as revised in 2008, and received approval from the University of Nottingham Faculty of Psychology Ethics Committee (reference: F1298R, 28/03/2022). After providing detailed information and answering all questions, participants provided written consent. The study was conducted in two sessions. During the first session, participants underwent a 45-minute MRI scan, which included a 14-minute resting-state fMRI sequence. They returned for a second session on a different day, scheduled at the same time of day as their initial visit (time difference: 55.4±40.1 minutes for the IFC group vs. 69.5±48.9 minutes for the thalamus group; p=0.469). Participants were pseudo-randomly assigned to one of two groups, ensuring an equal distribution of sexes, based on the TUS brain target: either the right inferior frontal cortex or the right thalamus. Immediately following stimulation, participants underwent another 45-minute MRI session (delay between TUS and rs-fMRI: 15±2.16 minutes for the IFC group vs. 15.4±1.37 minutes for the thalamus group; p=0.95), which included a 42-minute rs-fMRI sequence.

#### Data acquisition

During both sessions, MRI scans were conducted using a General Electric 3 Tesla scanner equipped with a 48-channel head coil. For the first session: A T1-weighted magnetization-prepared rapid gradient echo (MPRAGE) sequence was performed with a repetition time (TR) of 2.282 seconds, an echo time (TE) of 2.96 milliseconds, an inversion time (TI) of 800 milliseconds, a flip angle (FA) of 8°, and a field of view (FOV) of 256 × 256 mm, covering 180 slices with $1mm^3$ isotropic voxels. A Zero Echo Time (ZTE) sequence was also acquired with a TR of 0.531 seconds, a TE of 0.016 milliseconds, an FOV of 256 x 256 mm, and 176 slices with voxel dimensions of 1.016 x 1.016 x 1 $mm^3$. Shimming was performed using two echoes with a TE of 4.54 milliseconds, an FOV of 240 x 240 mm, and 32 slices with voxel dimensions of 3.75 x 3.75 x 5.8 $mm^3$ in the right-to-left direction. Additionally, a Diffusion-Weighted Imaging (DWI) sequence was acquired with a TR of 4.6 seconds, a TE of 90 milliseconds, 63 slices with voxel dimensions of 2.019 x 2.09 x 2 $mm^3$, in the right-to-left direction, and b-values of 0, 300, 1000, and 2000 $s/mm^2$ with 0, 10, 50, and 50 directions, respectively. A 14-minute resting-state functional MRI (rs-fMRI) sequence was also obtained with a TR of 1.4 seconds, a TE of 35 milliseconds, an FA of 68°, a FOV of 212 × 212 mm, and 88 interleaved slices with $2mm^3$ isotropic voxels and no slice gap, using a multiband acceleration factor of 3. Cardiac and respiratory data were recorded during this sequence, which was performed with the participants' eyes open.

For the second session: A 42-minute resting-state functional MRI sequence was conducted with the same parameters as the initial visit—TR of 1.4 seconds, TE of 35 milliseconds, FA of 68°, FOV of 212 × 212 mm, 88 interleaved slices with $2mm^3$ isotropic voxels, no slice gap, and a multiband acceleration factor of 3—along with cardiac and respiratory data registration. Additionally, a T1-weighted magnetization-prepared rapid gradient echo (MPRAGE) sequence was repeated with the same parameters as during the first visit: TR of 2.282 seconds, TE of 2.96 milliseconds, TI of 800 milliseconds, FA of 8°, and FOV of 256 × 256 mm, covering 180 slices with $1mm^3$ isotropic voxels.

#### Ultrasound stimulation

We applied the stimulation using a NeuroFUS PRO TPO-203 with the four-element CTX-500-4CH transducer (Sonic Concepts, Brainbox Ltd., Cardiff, United Kingdom). The theta-burst TUS protocol was used [2,3] as follows: central frequency=500kHz, pulse duration=20ms, pulse repetition interval=200ms (i.e., duty cycle=10%), total duration=80sec. The ISPPA was set at $54.51 W/cm^2$, following the safety guidelines [1]. To ensure effective coupling between the ultrasound transducer and the participant's head, ultrasound transmission gel was applied directly to the head, followed by application to the transducer face, and careful manual removal of any air bubbles.

#### fMRI pre-processing

The preprocessing pipeline consists of first removing signals from the ventricles, white matter (removed with Freesurfer), and cardiac and respiratory artifacts. Then, we despike the data and applied the afni_proc.py script, including the following steps: blocks (discarding the first two volumes), tshift, align (with the minimum outlier volume), tlrc, volreg, mask, scale, regress (including motion). Finally, we applied a 4mm smooth. The time-series were parcellated into 84 regions using the Desikan-Killiany atlas, including subcortical areas (see Supplementary Tables 2 and 3 for details).

#### Diffusion MRI pre-processing

DTI data were preprocessed using the standard approach provided by MRtrix, including denoising, removal of Gibbs artifacts, FSL preprocessing, B1 field inhomogeneity correction, and unsupervised estimation of brain tissues' multi-shell multi-tissue fiber orientation distributions. Subsequently, ten million tracts were estimated using a probabilistic approach before being reduced to 1 million after tcksift correction for streamline densities.

Analogous to the time-series, the DTI matrices were parcellated using the Desikan-Killiany atlas, including subcortical areas (see Supplementary Tables 2 and 3 for details), resulting in matrices with dimensions $\mathbb{R}^{84\times 84}$. This procedure derived the structural connectivity, measured as the average number of streamlines, and the distance, measured as the average length of streamlines between two regions. Three images were excluded due to excessive motion.

Table 1: Relative redundancy and synergy changes after TUS

| HOI changes | label | Lobe | t-score | p-value |
|---|---|---|---|---|
| **TUS-IFC** | | | | |
| redundancy | parsorbitalis L | frontal | 2.616 | 0.016 |
| redundancy | parsorbitalis R | frontal | 3.370 | 0.023 |
| redundancy | rostral middle frontal R | frontal | -2.448 | 0.046 |
| redundancy | paracentral R | frontal | -2.584 | 0.019 |
| redundancy | caudal middle frontal R | frontal | -2.093 | 0.046 |
| redundancy | caudate L | basal ganglia | -2.85 | 0.033 |
| redundancy | accumbens L | basal ganglia | -2.859 | 0.008 |
| synergy | rostral middle frontal L | frontal | -2.529 | 0.020 |
| synergy | rostral middle frontal R | frontal | -2.175 | 0.041 |
| synergy | supramarginal L | parietal | -2.125 | 0.048 |
| synergy | temporal pole L | temporal | -2.214 | 0.033 |
| synergy | entorhinal L | temporal | -2.111 | 0.039 |
| synergy | putamen L | basal ganglia | 2.755 | 0.015 |
| **TUS-Thal** | | | | |
| redundancy | parstriangularis L | frontal | 2.411 | 0.028 |
| redundancy | orbitofrontal R | frontal | 2.121 | 0.049 |
| redundancy | superior temporal L | temporal | 2.537 | 0.019 |
| redundancy | middle temporal R | temporal | -2.433 | 0.025 |
| redundancy | accumbens L | basal ganglia | 2.578 | 0.021 |
| redundancy | posterior cingulate R | cingulate | 3.478 | 0.003 |
| redundancy | lateral occipital R | occipital | -2.725 | 0.013 |
| synergy | rostral anterior L | cingulate | -2.125 | 0.048 |
| synergy | entorhinal L | temporal | -2.283 | 0.031 |
| synergy | pallidum L | basal ganglia | -3.061 | 0.006 |
| synergy | thalamus R | basal ganglia | 3.030 | 0.006 |

Table 2: Desikan-Killiany atlas, including subcortical areas: left hemisphere

| id | label | hemisphere | structure | Lobe |
|---|---|---|---|---|
| 1 | lateralorbitofrontal | L | cortex | frontal |
| 2 | medialorbitofrontal | L | cortex | frontal |
| 3 | frontalpole | L | cortex | frontal |
| 4 | parsorbitalis | L | cortex | frontal |
| 5 | parstriangularis | L | cortex | frontal |
| 6 | parsopercularis | L | cortex | frontal |
| 7 | rostralmiddlefrontal | L | cortex | frontal |
| 8 | caudalmiddlefrontal | L | cortex | frontal |
| 9 | superiorfrontal | L | cortex | frontal |
| 10 | precentral | L | cortex | frontal |
| 11 | paracentral | L | cortex | frontal |
| 12 | postcentral | L | cortex | parietal |
| 13 | superiorparietal | L | cortex | parietal |
| 14 | precuneus | L | cortex | parietal |
| 15 | inferiorparietal | L | cortex | parietal |
| 16 | supramarginal | L | cortex | parietal |
| 17 | rostralanteriorcingulate | L | cortex | cingulate |
| 18 | caudalanteriorcingulate | L | cortex | cingulate |
| 19 | posteriorcingulate | L | cortex | cingulate |
| 20 | isthmuscingulate | L | cortex | cingulate |
| 21 | temporalpole | L | cortex | temporal |
| 22 | inferiortemporal | L | cortex | temporal |
| 23 | middletemporal | L | cortex | temporal |
| 24 | superiortemporal | L | cortex | temporal |
| 25 | transversetemporal | L | cortex | temporal |
| 26 | bankssts | L | cortex | temporal |
| 27 | fusiform | L | cortex | temporal |
| 28 | entorhinal | L | cortex | temporal |
| 29 | parahippocampal | L | cortex | temporal |
| 30 | insula | L | cortex | temporal |
| 31 | cuneus | L | cortex | occipital |
| 32 | lingual | L | cortex | occipital |
| 33 | pericalcarine | L | cortex | occipital |
| 34 | lateraloccipital | L | cortex | occipital |
| 35 | hippocampus | L | subcortex/brainstem | basal |
| 36 | amygdala | L | subcortex/brainstem | basal |
| 37 | thalamusproper | L | subcortex/brainstem | basal |
| 38 | accumbensarea | L | subcortex/brainstem | basal |
| 39 | caudate | L | subcortex/brainstem | basal |
| 40 | putamen | L | subcortex/brainstem | basal |
| 41 | pallidum | L | subcortex/brainstem | basal |
| 42 | brainstem | L | subcortex/brainstem | brainstem |

Table 3: Desikan-Killiany atlas, including subcortical areas: right hemisphere

| id | label | hemisphere | structure | Lobe |
|---|---|---|---|---|
| 43 | lateralorbitofrontal | R | cortex | frontal |
| 44 | medialorbitofrontal | R | cortex | frontal |
| 45 | frontalpole | R | cortex | frontal |
| 46 | parsorbitalis | R | cortex | frontal |
| 47 | parstriangularis | R | cortex | frontal |
| 48 | parsopercularis | R | cortex | frontal |
| 49 | rostralmiddlefrontal | R | cortex | frontal |
| 50 | caudalmiddlefrontal | R | cortex | frontal |
| 51 | superiorfrontal | R | cortex | frontal |
| 52 | precentral | R | cortex | frontal |
| 53 | paracentral | R | cortex | frontal |
| 54 | postcentral | R | cortex | parietal |
| 55 | superiorparietal | R | cortex | parietal |
| 56 | precuneus | R | cortex | parietal |
| 57 | inferiorparietal | R | cortex | parietal |
| 58 | supramarginal | R | cortex | parietal |
| 59 | rostralanteriorcingulate | R | cortex | cingulate |
| 60 | caudalanteriorcingulate | R | cortex | cingulate |
| 61 | posteriorcingulate | R | cortex | cingulate |
| 62 | isthmuscingulate | R | cortex | cingulate |
| 63 | temporalpole | R | cortex | temporal |
| 64 | inferiortemporal | R | cortex | temporal |
| 65 | middletemporal | R | cortex | temporal |
| 66 | superiortemporal | R | cortex | temporal |
| 67 | transversetemporal | R | cortex | temporal |
| 68 | bankssts | R | cortex | temporal |
| 69 | fusiform | R | cortex | temporal |
| 70 | entorhinal | R | cortex | temporal |
| 71 | parahippocampal | R | cortex | temporal |
| 72 | insula | R | cortex | temporal |
| 73 | cuneus | R | cortex | occipital |
| 74 | lingual | R | cortex | occipital |
| 75 | pericalcarine | R | cortex | occipital |
| 76 | lateraloccipital | R | cortex | occipital |
| 77 | hippocampus | R | subcortex/brainstem | basal |
| 78 | amygdala | R | subcortex/brainstem | basal |
| 79 | thalamusproper | R | subcortex/brainstem | basal |
| 80 | accumbensarea | R | subcortex/brainstem | basal |
| 81 | caudate | R | subcortex/brainstem | basal |
| 82 | putamen | R | subcortex/brainstem | basal |
| 83 | pallidum | R | subcortex/brainstem | basal |
| 84 | brainstem | R | subcortex/brainstem | brainstem |

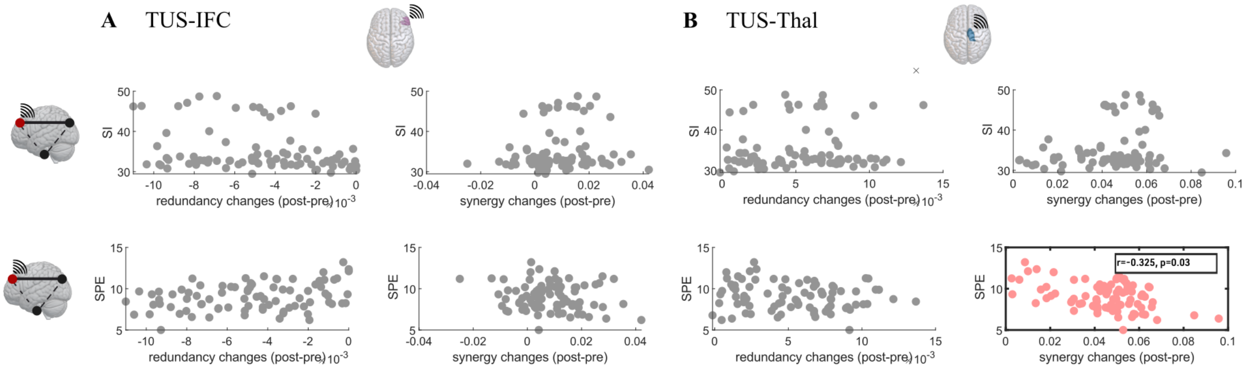


Figure 1: For TUS-IFC **A.** and TUS-Thal **B.** Within each subpanel, the rows correspond to the search information (SI) and short path efficiency (SPE) models, while each column to changes in informational quantities (redundancy, left column; synergy, left right column). The darker boxes represent the p-values lower than 0.05 after a Bonferroni correction.

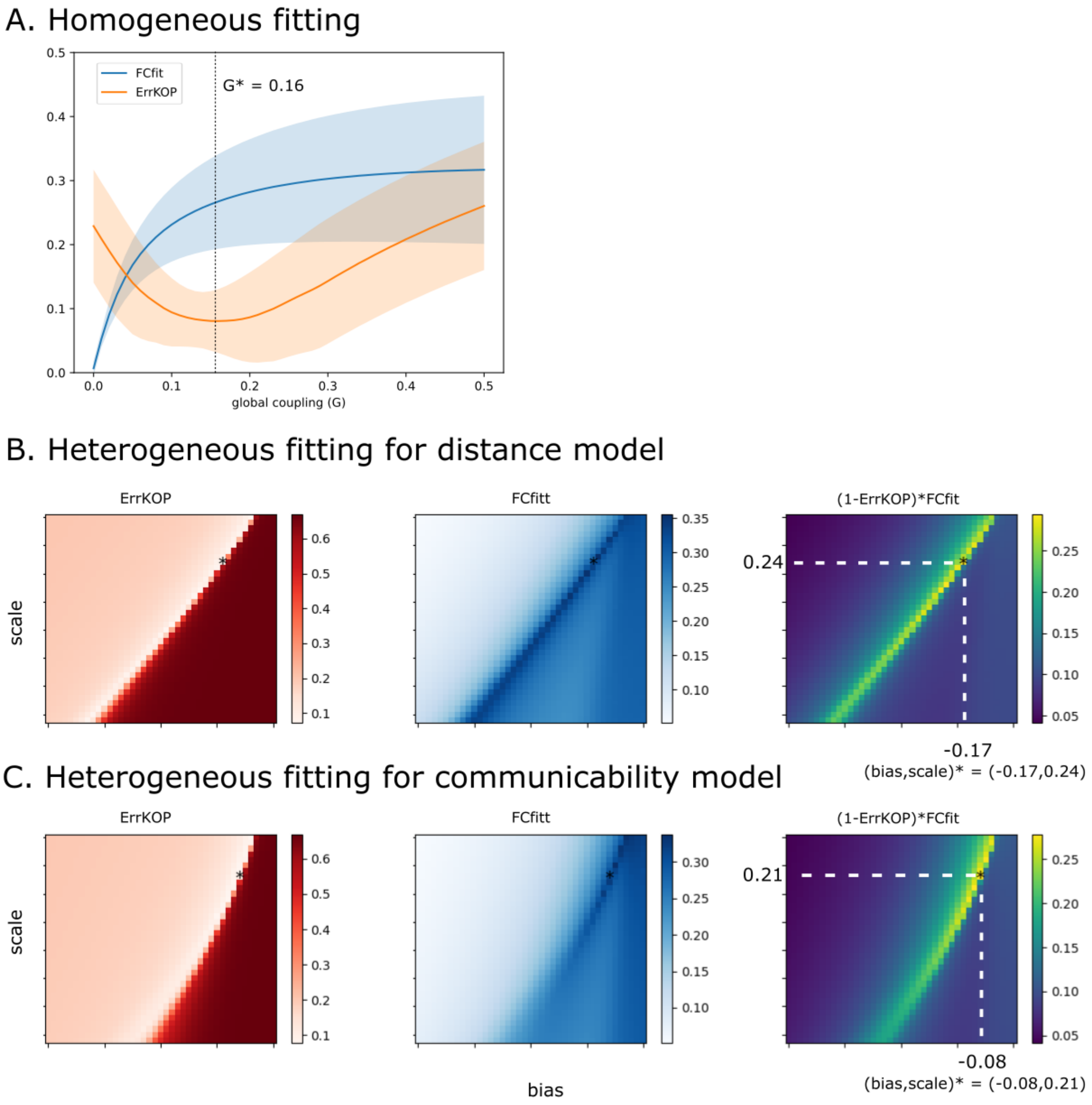


Figure 2: **A**. Homogeneous model for the control condition (non-TUS) **B.** Heterogeneous distance model for non-TUS. **C.** Heterogeneous communicability model for non-TUS. The first column corresponds to the differences in synchrony (measured as the mean of the Kuramoto order parameter) between the empirical and simulated data. The second column is the correlation between the functional connectivities between the simulations and the empirical data, and the third column is the multiplication of the two former columns.

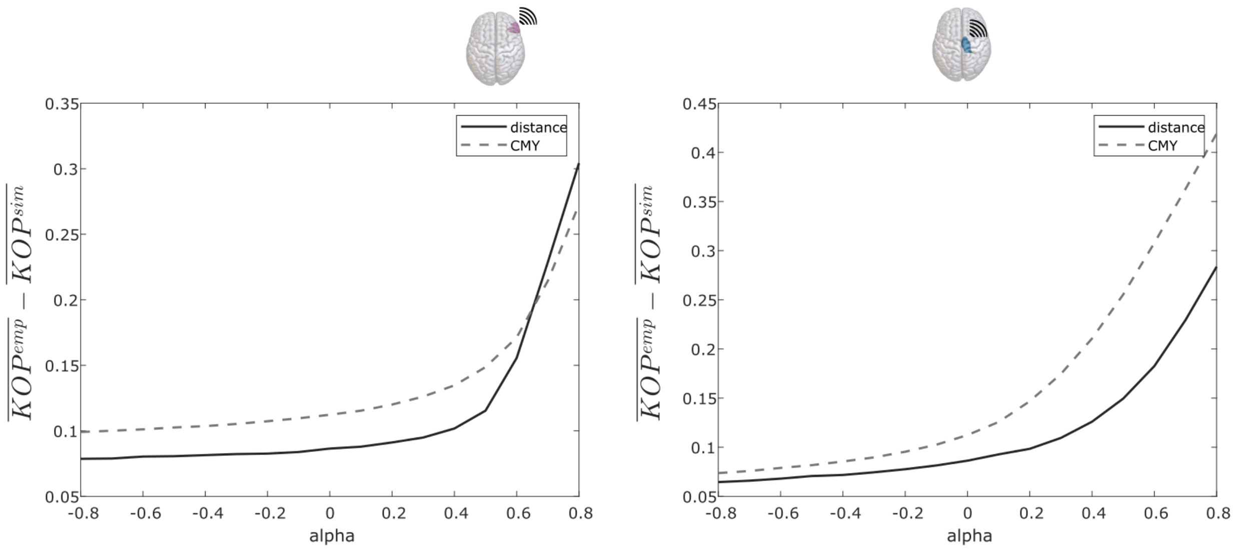


Figure 3: Synchrony differences (mean(KOP)) between empirical data and the target at various intensities, modulated by $\alpha$, for each model (distance in solid line, and communicability in dashed line).

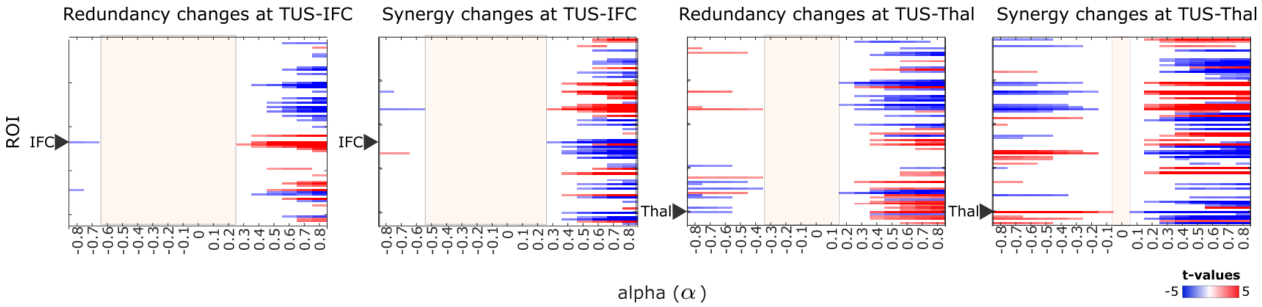


Figure 4: Corrected t-values in the simulated data (TUS minus control), for the model based on communicability. We displayed the significant t-values corrected using a permutation test with N = 1000 iterations.

### Bibliography


[1] Jean-Francois Aubry, David Attali, Mark Schafer, Elsa Fouragnan, Charles Caskey, Robert Chen, Ghazaleh Darmani, Ellen J Bubrick, Jérôme Sallet, Christopher Butler, Charlotte Stagg, Miriam Klein-Flügge, Seung-Schik Yoo, Brad Treeby, Lennart Verhagen, and Kim Butts Pauly. Itrusst consensus on biophysical safety for transcranial ultrasonic stimulation.

[2] Siti N. Yaakub, Tristan A. White, Jamie Roberts, Eleanor Martin, Lennart Verhagen, Charlotte J. Stagg, Stephen Hall, and Elsa F. Fouragnan. Transcranial focused ultrasound-mediated neurochemical and functional connectivity changes in deep cortical regions in humans. *Nature Communications*, 14:5318, 9 2023.

[3] Ke Zeng, Ghazaleh Darmani, Anton Fomenko, Xue Xia, Stephanie Tran, Jean François Nankoo, Yazan Shamli Oghli, Yanqiu Wang, Andres M. Lozano, and Robert Chen. Induction of human motor cortex plasticity by theta burst transcranial ultrasound stimulation. *Annals of Neurology*, 91:238–252, 2 2022.
